# The functional divergence of two ethylene receptor subfamilies that exhibit Ca^2+^-permeable channel activity

**DOI:** 10.1101/2025.11.28.691086

**Authors:** Chenliang Pan, Junyuan Cheng, Zining Lin, Dongdong Hao, Zhina Xiao, Yuhang Ming, Wen Song, Li Liu, Hongwei Guo

## Abstract

Ethylene is a gaseous plant hormone crucial for regulating plant growth, development, and stress adaptation, yet the molecular basis underlying ethylene receptor function remains elusive. Here, we show that subfamily I receptors constitute core ethylene-sensing module and function epistatically to subfamily II receptors. Notably, we discover that only subfamily I receptors possess Ca^2+^-permeable channel activity, which are indispensable for ethylene-induced cytosolic calcium influx. Overall, this work supports a mechanistic framework in which subfamily I receptors integrate ethylene sensing with Ca^2+^ influx, providing new insight into how plants translate hormonal cues into downstream signaling events.

## Introduction

As one of the earliest identified phytohormones, ethylene orchestrates a wide array of plant development processes and adaptive responses to environmental stress (Schaller and Kieber, 2002). Foundational insights into ethylene biology were driven by forward genetic screens in *Arabidopsis thaliana*, which exploit the characteristic “triple response” phenotype—exaggerated apical hook curvature, shortened hypocotyl and root, and radial swelling of the hypocotyl—to isolate ethylene-response mutants (Guzman and Ecker, 1990). Nearly four decades of research have since led to the establishment of a signaling pathway that connects ethylene perception at the endoplasmic reticulum (ER) membrane to transcriptional reprograming in the nucleus (Hao et al., 2025). Despite substantial progress in elucidating downstream signaling events in recent years, the molecular mechanism by which the ethylene receptors at the apex of this pathway perceive the gaseous hormone remains a fundamental and unresolved question in the field (Binder, 2020; Hao *et al*., 2025).

In *Arabidopsis*, the ethylene receptor family consists of five members: ETHYLENE RESPONSE 1 (ETR1), ETHYLENE RESPONSE SENSOR (ERS1), ETHYLENE INSENSITIVE 4 (EIN4), ETR2 and ERS2 (Bleecker et al., 1988; Chang et al., 1993; Hua et al., 1995; Hua et al., 1998; Roman et al., 1995; Sakai et al., 1998). They are ER-localized transmembrane proteins, negatively regulating the ethylene signal transduction (Chen et al., 2002; Hua and Meyerowitz, 1998). Based on the presence or absence of a conserved histidine kinase motif, they are classified into two subfamilies: subfamily I receptors, ETR1 and ERS1, retain the conserved histidine kinase motif, whereas subfamily II receptors, EIN4, ETR2 and ERS2, lack this motif (Azhar et al., 2019; Hao *et al*., 2025). Structurally, subfamily II receptors also differ from subfamily I receptors by the presence of an additional N-terminal region, which has been proposed to represent an extra transmembrane helix (Azhar *et al*., 2019; Binder, 2020). In addition, genetic analyses have highlighted profound functional differences between the two subfamilies. Double loss-of-function mutants in the subfamily I receptors exhibit a dramatic constitutive ethylene-response phenotype culminating in postembryonic lethality—a phenotype far more severe than that of the subfamily II triple mutant (Hua and Meyerowitz, 1998; Qu et al., 2007). From a dynamic perspective, gain-of-function mutations in subfamily I receptors abolish ethylene responsiveness in both the rapid growth-inhibition phase (phase I) and the transcription-dependent phase (phase II) (Azhar et al., 2023; Chen et al., 2019). By contrast, gain-of-function mutations in subfamily II receptors permit nearly normal phase I and phase II responses, followed by gradually resuming the growth rate of untreated plants after phase II (Chen *et al*., 2019). Together, these findings suggest that the two receptor subfamilies might act through distinct mechanisms. Notably, the kinase activity of ethylene receptors has shown to be dispensable for ethylene signaling (Chen et al., 2009; Hall et al., 2012), leaving the biochemical basis of receptor function poorly defined. Thus, although ethylene receptors were the first genetically-identified plant hormone receptors (Bleecker et al., 1988; Chang et al., 1993), their mechanism of action remains among the least understood.

Calcium is a unique mineral nutrient that also serves as a ubiquitous signaling molecule in all eukaryotes (Luan and Wang, 2021). In plants, Ca^2+^ functions as a dynamic intracellular messenger that participates in virtually all developmental and stress-related signaling pathways (Reddy et al., 2011). Previous studies have revealed a potential interplay between Ca^2+^ signaling and ethylene response. For instance, early work demonstrated that the Ca^2+^ chelator EGTA could block ethylene-dependent induction of chitinase (Raz and Fluhr, 1992). Subsequent study showed that ethylene promoted Ca^2+^ influx into excised pea stem segments (Kwak and Lee, 1997). In addition, ethylene was reported to activate unidentified plasma membrane Ca^2+^-permeable channels in tobacco suspension cells (Zhao et al., 2007). Despite these findings, the direct link between Ca^2+^ and ethylene signaling remains largely unknown.

In this study, we dissected the functional hierarchy and biochemical properties of the two ethylene receptor subfamilies. We showed that plants lacking both subfamily I receptors were completely unresponsive to ethylene, establishing that subfamily I receptors constitute the core ethylene-sensing module in *Arabidopsis*. Genetic analyses further demonstrated that the five receptors do not occupy equivalent positions within the signaling pathway, and that subfamily II receptors require subfamily I receptors to function. Structural predictions indicated that ethylene receptors may operate as ion channels. Consistent with this, electrophysiological analyses revealed that subfamily I, but not subfamily II, receptors exhibited Ca^2+^-permeable channel activity. Moreover, we showed that ethylene elicits Ca^2+^ influx in a manner dependent exclusively on subfamily I receptors. Together, these findings uncover a previously unrecognized role of ethylene receptors as Ca^2+^ channels, thereby directly linking Ca^2+^ signaling to ethylene perception.

## Results

### Subfamily I receptors are indispensable for ethylene perception in *Arabidopsis*

Previous study showed that the subfamily I receptor loss-of-function double mutant *etr1-7 ers1-3* displayed an extreme constitutive triple-response phenotype and post-embryonic lethality (Qu *et al*., 2007), which is not suitable for further investigating ethylene-induced phenotypic alterations. Notably, the developmental defects of *etr1-7 ers1-2* mutants could be fully suppressed by the mutation of EIN2, a downstream core positive component (Alonso et al., 1999; Hall and Bleecker, 2003). To further delineate the function of the subfamily I receptors in ethylene perception, we introduced an β-estradiol inducible *EIN2* transgene into the *etr1-7 ers1-3 ein2-5* background (*iEIN2-CFP-HA/etr1-7 ers1-3 ein2-5,* hereafter *iEIN2/11ein2*) (Supplemental Figure 1 and 2). This transgenic line can bypass the post-embryonic lethality in the absence of β-estradiol, while preserve the integrity of the downstream ethylene signaling in the presence of β-estradiol, thereby providing a tractable system for dissecting the roles of two subfamily I receptors in ethylene responses. To enable direct comparison between the two receptor subfamilies, we generated two additional lines: *iEIN2-CFP-HA/etr2-3 ers2-3 ein4-4 ein2-5* (hereafter *iEIN2/224ein2*) that lacks three subfamily II receptors, and *iEIN2-CFP-HA/etr1-7 ers1-3 etr2-3 ers2-3 ein4-4 ein2-5* (hereafter *iEIN2/11224ein2*) that lacks all five ethylene receptors.

Seedlings lacking the two subfamily I receptors, *iEIN2/11ein2*, displayed a strong constitutive ethylene response phenotype on MS medium containing 10 μM β-estradiol, which resembled the severe phenotype of *etr1-7 ers1-3* (Qu *et al*., 2007). This phenotype was also indistinguishable from that of plants lacking all five receptors, *iEIN2/11224ein2,* upon β-estradiol treatment (Figure 1A). Furthermore, in both *iEIN2/11ein2* and *iEIN2/11224ein2*, application of 10 μM ethylene biosynthetic precursor 1-aminocyclopropane-1-carboxylic acid (ACC) failed to elicit any further response (Figure 1A). By contrast, plants lacking all subfamily II receptors, *iEIN2/224ein2*, showed a modest constitutive ethylene-response phenotype and remained responsive to ACC (Figure 1A). Because the constitutive ethylene-response phenotypes induced by 10 μM β-estradiol were extremely strong and might mask potential ethylene responses, we then titrated the inducer concentrations. We found that *iEIN2/11ein2* and *iEIN2/11224ein2* seedlings failed to respond to ethylene at any β-estradiol concentration between 8 and 1000 nM, whereas *iEIN2/ein2* responded evidently across the same concentration range (Figure 1B and 1C). *iEIN2/224ein2* displayed a weak constitutive ethylene-response phenotype in the air (Figure 1A). Once the β-estradiol concentration exceeded 40 nM, this constitutive phenotype was no longer intensified (Figure 1B and 1C), supporting that loss of subfamily II receptors leads to partial activation of the downstream signaling. Together, these results demonstrate that subfamily I, but not subfamily II, receptors are indispensable for the ethylene-induced triple response.

**Figure 1.**
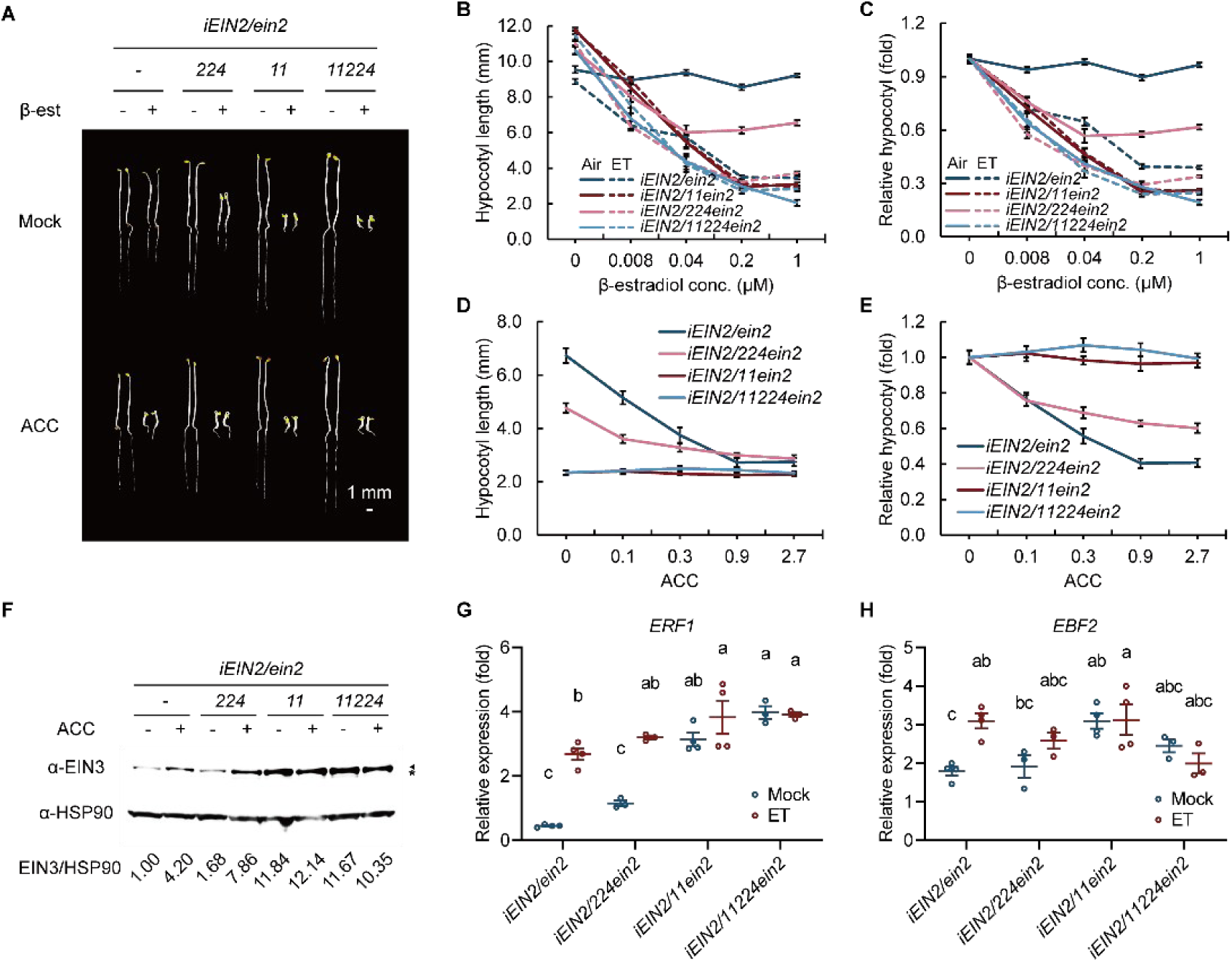
Subfamily I receptors are required for ethylene perception in *Arabidopsis*. (A) Triple response phenotypes of *iEIN2/ein2-5* (*iEIN2/ein2*), *iEIN2/etr2-3 ers2-3 ein4-4 ein2-5* (*iEIN2/224ein2*), *iEIN2/etr1-7 ers1-3 ein2-5* (*iEIN2/11ein2*), and *iEIN2/etr1-7 ers1-3 etr2-3 ers2-3 ein4-4 ein2-5* (*iEIN2/11224ein2*) plants. Seedlings were grown on MS medium supplemented with or without 10 μM ACC in the dark for 3.5 d (Scale bar, 1 mm.). (B) Quantification of hypocotyl length in 3.5-day-old etiolated seedlings. Solid lines indicate air-treated samples, and dashed lines indicate ethylene-treated samples. Seedlings were grown on MS medium supplemented with the indicated concentrations of β-estradiol. (C) Normalized hypocotyl length based on the data shown in (B). Values represent means ± SE (n ≥ 12) in (B) and (C). (D) Quantification of hypocotyl length in etiolated seedlings 48 h after germination through dynamic platform imaging. Seedlings were grown on MS medium containing 20 nM β-estradiol and the indicated concentrations of ACC (μM). (E) Normalized hypocotyl length based on the data shown in (D). Values represent means ± SE (n ≥ 15) in (D) and (E). (F) Immunoblot assays showing EIN3 protein levels. Triangles indicate the EIN3 bands, and the star symbol marks a nonspecific band. HSP90 was used as a loading control. Seedlings were grown on ½ MS medium with or without ACC at 22 °C for 7 days, sprayed with 10 μM β-estradiol, and harvested 4 h later. (G–H) Relative transcript levels of *ERF1* (G) and *EBF1* (H) in seedlings of the indicated genotypes. Seedlings were grown on ½ MS medium at 22 °C for 7 days, sprayed with 10 μM β-estradiol, and after 8 h exposed to either air or ethylene. Values represent means ± SE (n ≥ 3). Statistical significances between samples were analyzed by two-way ANOVA and indicated by different lowercase letters for *P* ≤ 0.05.

Since the ACC dose used above was nearly saturating, we also examined the responses of those lines to a gradient of ACC concentrations upon a modest β-estradiol application (20 nM). Consistently, *iEIN2/11ein2* seedlings showed no ethylene response at any tested ACC concentrations (Figure 1D and 1E), reinforcing the conclusion that ethylene responses in *Arabidopsis* strictly depend on subfamily I receptors. Compared with *iEIN2/ein2*, *iEIN2/224ein2* seedlings exhibited a saturated ethylene-response phenotype at relatively low ACC concentrations (Figure 1D and 1E), suggesting that loss of subfamily II receptors leads to the maturation of ethylene perception at a lower hormone dosage. Taken together, these results indicate that subfamily I receptors are indispensable for the ethylene perception at any hormone dosage, while subfamily II receptors mainly contribute to ethylene perception in a higher dosage range.

To further investigate whether plants lacking subfamily I ethylene receptors are still capable of mounting transient ethylene responses, we examined EIN3 protein levels, a core transcription factor whose protein accumulates in response to ethylene (Gagne et al., 2004; Guo and Ecker, 2003; Potuschak et al., 2003), in the different receptor mutants with or without ACC treatments. In the presence of 10 μM β-estradiol, EIN3 protein accumulated in response to ACC in both *iEIN2/ein2* and *iEIN2/224ein2*, whereas *iEIN2/11ein2* and *iEIN2/11224ein2* showed constitutively elevated EIN3 levels, and did not increase further upon ACC treatment (Figure 1F). Consistently, the ethylene-responsive genes *ETHYLENE RESPONSE FACTOR 1* (*ERF1)* and *EIN3-BINDING F BOX PROTEIN 2 (EBF2)* were constitutively upregulated and lost their response to ethylene in *iEIN2/11ein2* and *iEIN2/11224ein2* (Figure 1G and 1H). These molecular data further support that subfamily I receptors are essential for ethylene signal transduction.

### Subfamily I receptors are genetically epistatic to subfamily II receptors

Previous studies showed that all ethylene receptors can bind ethylene (O’Malley et al., 2005), and that both subfamily I and subfamily II receptors possess gain-of-function mutations that confer nearly complete ethylene insensitivity (Bleecker *et al*., 1988; Chang *et al*., 1993; Hua *et al*., 1995; Hua and Meyerowitz, 1998; Hua *et al*., 1998; Roman *et al*., 1995). Therefore, the two receptor subfamilies are generally considered to act in parallel in the ethylene signaling pathway, despite differences in the relative strength of their functional contributions (Binder, 2020; Qu *et al*., 2007). However, our results above indicated that only subfamily I receptors are essential for ethylene responses, prompting us to reconsider the genetic relationship of the two receptor subfamilies in the ethylene signaling pathway. Given that the subfamily II gain-of-function mutants *ein4-1* and *etr2-1* are completely insensitive to ethylene, we introduced these mutations separately into the subfamily I double loss-of-function mutant *etr1-7 ers1-3* background. We found that *etr1-7 ers1-3* fully suppressed the phenotypes of *ein4-1* and *etr2-1*, resulting instead in the constitutive ethylene-response phenotype characteristic of *etr1-7 ers1-3* (Figure 2). In contrast, the transgenic gain-of-function mutation *ers1-1* could efficiently rescue the phenotype of *etr1-7 ers1-3* (Figure 2). These findings suggest that genetical functions of two receptor subfamilies are not in parallel, wherein subfamily I receptors are epistatic to subfamily II receptors.

**Figure 2.**
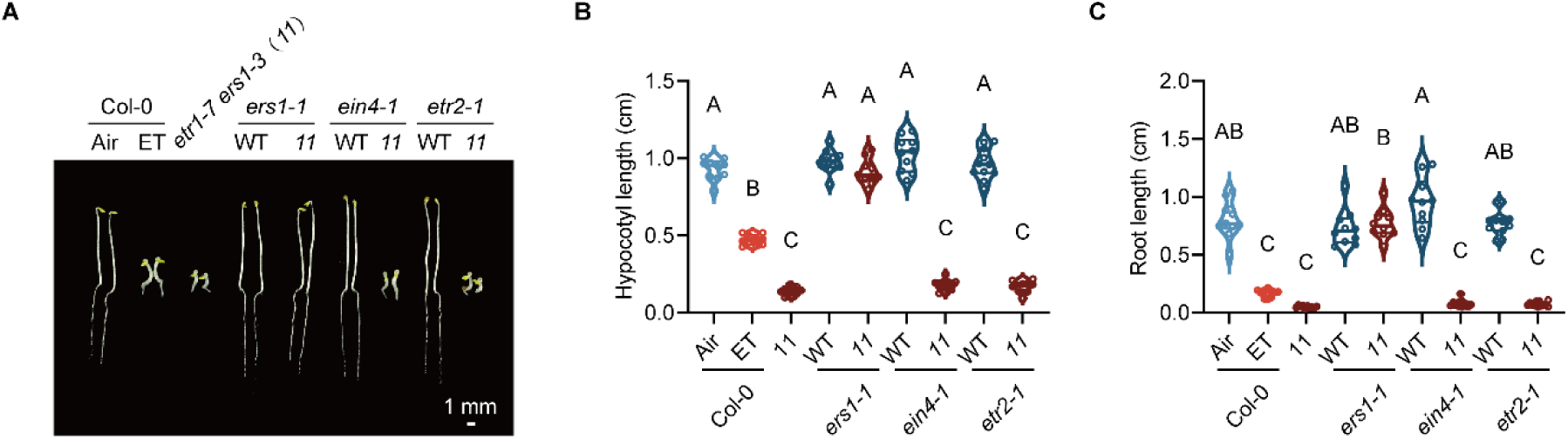
Subfamily I receptors are genetically epistatic to subfamily II receptors. (A) Triple response phenotypes of Col-0 (in air or ET), *etr1-7 ers1-3* (abbreviated as *11*), *ers1-1*, *ers1-1 etr1-7 ers1-3*, *ein4-1*, *ein4-1 etr1-7 ers1-3*, *etr2-1* and *etr2-1 etr1-7 ers1-3* plants. Seedlings were grown on MS medium in the dark for 3.5 d (Scale bar, 1 mm.). (B-C) Quantification of hypocotyl (B) and main root (C) length of 3.5-day-old etiolated seedlings in (A). In this experiment, the sample size (n) was greater than 10. Statistical significances between samples were analyzed by one-way ANOVA and indicated by different uppercase letters for *P* ≤ 0.01.

In addition, we found that 1-methylcyclopropene (1-MCP), a potent ethylene antagonist that can efficiently block ethylene response (Binder, 2020), could rescue the constitutive ethylene-response phenotype of *etr2-3 ers2-3 ein4-4* triple mutant, but not that of *etr1-7 ers1-3* (Supplemental Figure 3). Previous studies have shown that 1-MCP enhances receptor activity, rendering them in a constitutively active state (Schaller and Binder, 2017). In the presence of 1-MCP, the subfamily II receptors in *etr1-7 ers1-3* background are presumably in a gain-of-function state; however, they still failed to rescue the phenotype of *etr1-7 ers1-3* (Supplemental Figure 3). By contrast, 1-MCP-induced activation of subfamily I receptors effectively rescued the phenotype of *etr2-3 ers2-3 ein4-4* (Supplemental Figure 3). The different response to 1-MCP provides additional evidence to support that subfamily I receptors are epistatic to subfamily II receptors, implying potentially different biochemical properties of the two subfamily receptors.

### Subfamily I ethylene receptors ETR1 and ERS1 are Ca^2+^-permeable channels

The functional divergence of two subfamily ethylene receptors promoted us to explore the uncharacterized biochemical function of ethylene receptors. We first used AlphaFold3-based structural simulation for seeking potential molecular clues (Abramson et al., 2024). Since previous data indicated that ethylene receptors can form homodimers (Schaller et al., 1995), we simulated the homo-dimeric structures of the five ethylene receptors. The AlphaFold3 predictions showed that ethylene receptors form symmetric dimers (Supplemental Figure 4A). Notably, the AlphaFold3 structure of ERS1 dimer is highly aligned with the previously reported crystal structure of dimeric histidine phosphotransfer domains of ERS1 (Mayerhofer et al., 2015) (Supplemental Figure 4B), suggesting that AlphaFold3-based structural prediction matches the experimental observation. Intriguingly, the N-terminal transmembrane domains of two ETR1 protomers assemble into a channel-like structure (Supplemental Figure 4C). Six transmembrane helices form a pore with two-fold symmetry. The charged residues are located in the inner surface of α-2 and α-3 helices for potential ion selective permeation (Supplemental Figure 4C). Considering that ethylene receptors are located in the ER membrane (Chen *et al*., 2002), ethylene receptors could potentially function as ion channel permeating Ca^2+^ from ER to the cytoplasm. Thus, we further used AlphaFold3 to simulate Ca^2+^ interaction with ETR1. As expected, the Ca^2+^ ion binds to the inner cavity of the N-terminal transmembrane domain of ETR1 (Figure 3A).

**Figure 3.**
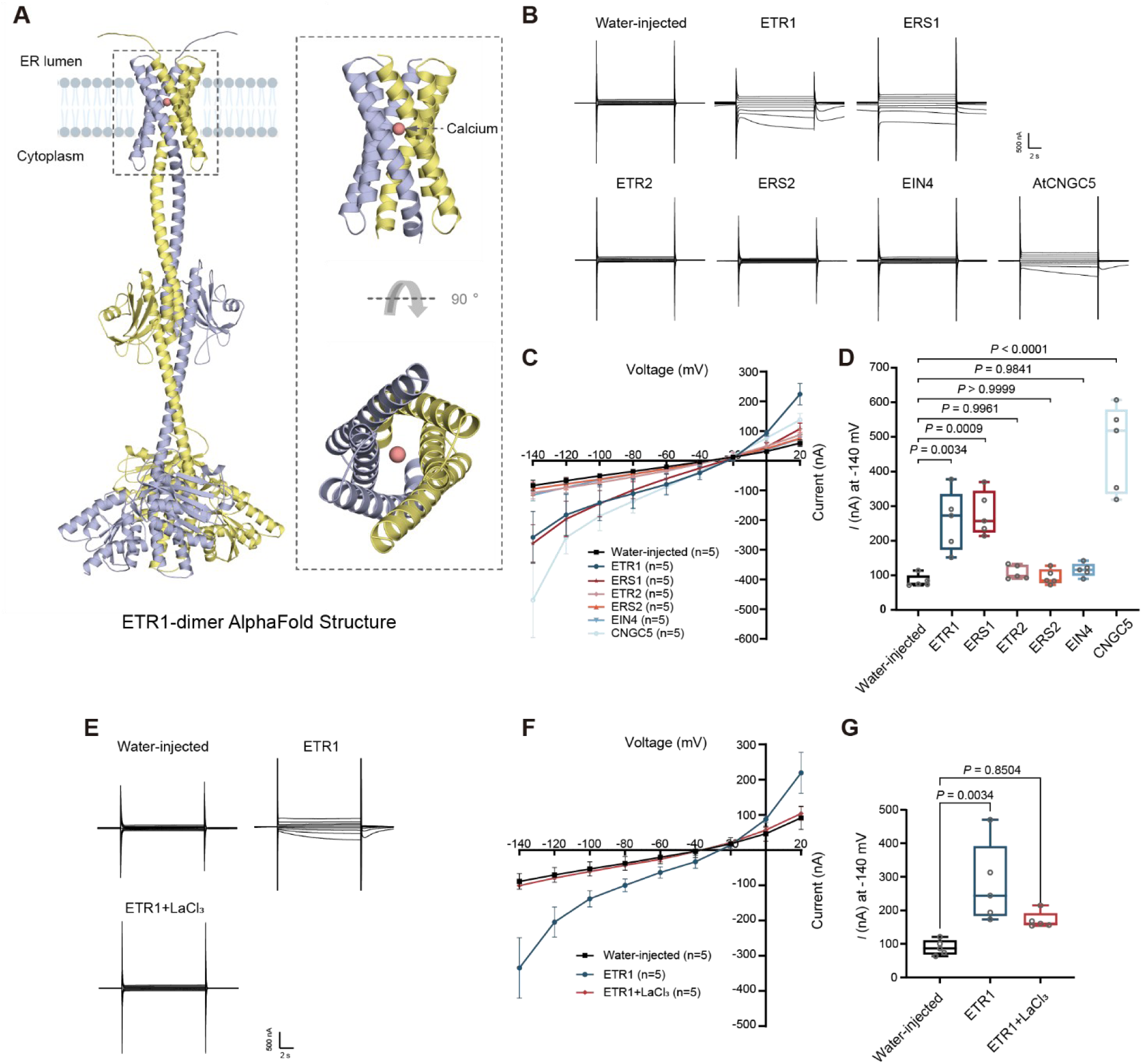
Subfamily I receptors ETR1 and ERS1 are Ca^2+^-permeable channels. (A) AlphaFold3 structural simulation of ETR1 dimer in complex with Calcium. Left panel, ERT1 protomers are shown in colors of light blue and yellow. The Calcium ion is shown in pink. The ER membrane is shown in cartoon. The N-terminal transmembrane domain of ETR1 dimer is highlighted with dot box. Right panel, zoom in the N-terminal transmembrane domains of ETR1 dimer with two angles. The Calcium ion is indicated with an arrow. (B) TEVC recordings from *Xenopus oocytes* expressing ETR1, ERS1, ETR2, ERS2 or EIN4 in bath solution containing 30 mM CaCl_2_. Water-injected was employed as a negative control, AtCNGC5 was employed as a positive control. (C) Steady-state I-V curves of multiple recordings of ETR1, ERS1, ETR2, ERS2, EIN4 or AtCNGC5. (D) Absolute values of current amplitudes at -140 mV from multiple recordings of ETR1, ERS1, ETR2, ERS2, EIN4 or AtCNGC5. (E) TEVC recordings from Xenopus oocytes expressing ETR1 in bath solution containing 30 mM CaCl_2_ or 1 mM LaCl_3_. (F) Steady-state I-V curves of multiple recordings of ETR1. (G) Absolute values of current amplitudes at -140 mV from multiple recordings of ETR1. Data are means ± SD (n=5). Different letters represent significant differences at *P* < 0.05 (one-way ANOVA and Tukey’s multiple comparison tests). Three biologically independent experiments with different batches of oocytes were carried out with similar results.

To investigate whether ethylene receptors function as Ca^2+^-permeable channels, we performed two-electrode voltage clamp (TEVC) electrophysiological assays in the *Xenopus oocytes* reconstitution system. Each of the five *Arabidopsis* ethylene receptors was expressed in *Xenopus oocytes* (Supplemental Figure 5A). Strikingly, TEVC recordings showed that oocytes expressing subfamily I receptor ETR1 and ERS1 produced pronounced inward Ca^2+^ currents upon hyperpolarization, which was similar to a positive control using Arabidopsis Ca^2+^-permeable cyclic nucleotide-gated channel 5 (CNGC5) (Figure 3B-D). In contrast, oocytes expressing subfamily II receptor ETR2, ERS2 and EIN4 did not produce detectable currents (Figure 3B-D). Adding a typical Ca^2+^ channel blocker LaCl_3_ completely inhibited Ca^2+^ permeable activity of ETR1 (Figure 3E-G). These results demonstrate that subfamily I, but not subfamily II, ethylene receptors possess Ca^2+^ permeable channel activity.

### Key residues in N-terminal domain of ethylene receptors impact Ca^2+^-permeability

Next, we tried to identify the key residues involved in discriminating channel activity of subfamily I and II ethylene receptors. Although the overall structures of subfamily I and II receptors are very similar (Supplemental Figure 4A), a key residue ETR1^F76^ located in the N-terminal inner cavity drew our attention (Figure 4A). Sequence alignments show that the Phe residue is highly conserved in subfamily I receptors, while Phe is substituted to Tyr in subfamily II receptors (Figure 4B). Structural analysis indicates that the substitution of Phe to Tyr substantially alters the polar interactions of charged residues in the ion channel selective region (Figure 4A). Therefore, we expressed and examined the channel activity of ETR1^F76Y^ substitution in *Xenopus oocytes* (Supplemental Figure 5B). The TEVC recordings showed that the Ca^2+^-permeable activity of ETR1^F76Y^ was largely compromised in comparison to that of ETR1^WT^ (Figure 4C-E). Thus, these results suggest that sequence polymorphism determines the Ca^2+^ permeable channel activity of the two subfamily ethylene receptors.

**Figure 4.**
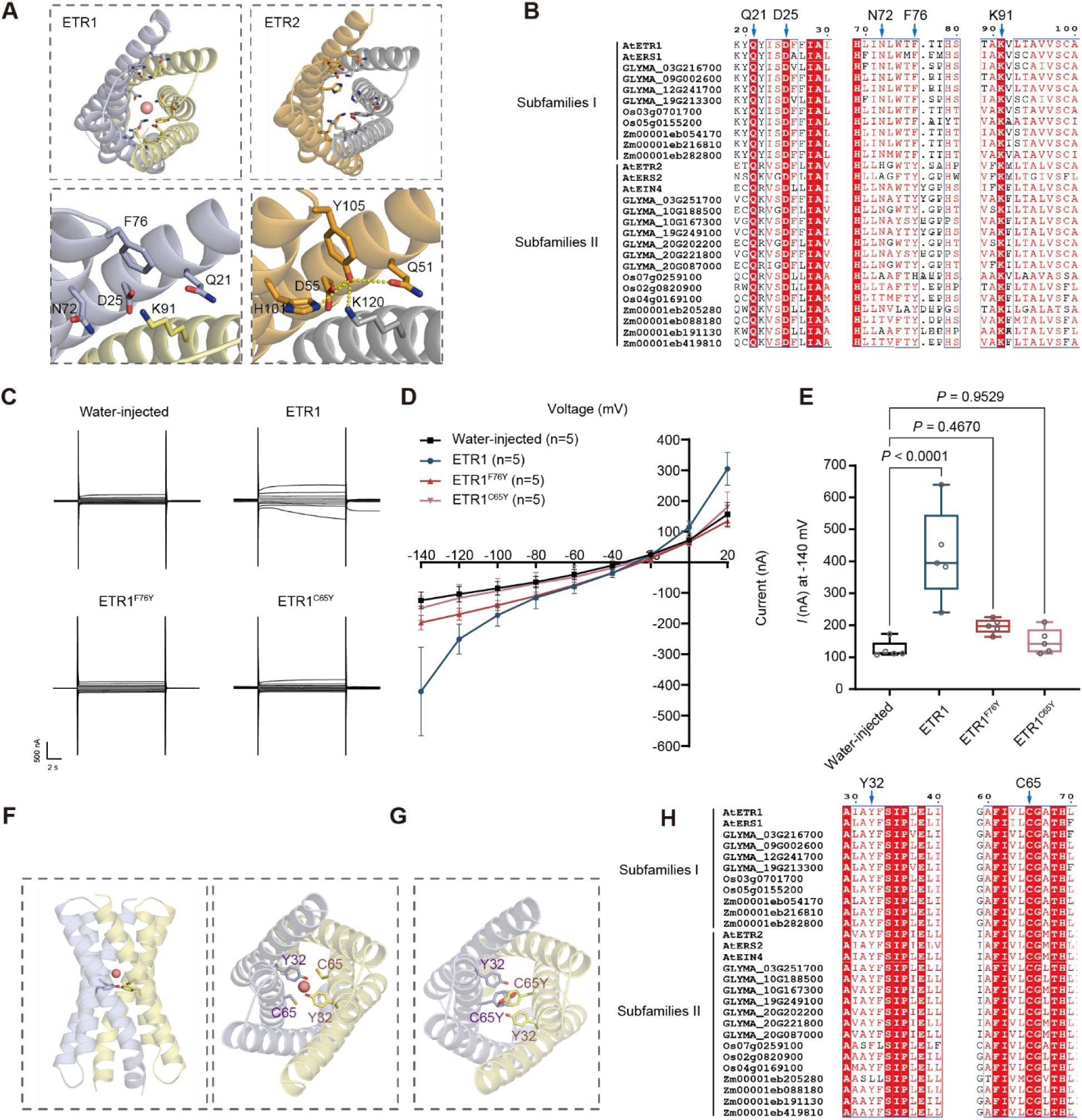
The key residues in the N-terminal domains of ethylene receptors impact Ca^2+^-permeable channel activity. (A) Structural analysis of charged residues in the N-terminal cavity of ETR1 and ETR2. Left, the charged resides of ETR1 are shown in sticks and labeled. Right, the charged resides of ETR2 are shown in sticks and labeled. The polar interactions are indicated with yellow dots. (B) and (H) Sequence alignments of ethylene receptors from different species. The subfamily I and II are labeled. The functional conserved residues are indicated with arrows. (C) TEVC recordings from *Xenopus oocytes* expressing ETR1, ETR1^F76Y^ or ETR1^C65Y^ in bath solution containing 30 mM CaCl_2_. (D) Steady-state I-V curves of multiple recordings of ETR1, ETR1^F76Y^ or ETR1^C65Y^. (E) Absolute values of current amplitudes at -140 mV from multiple recordings of ETR1, ETR1^F76Y^ or ETR1^C65Y^. Data are means ± SD (n=5). Different letters represent significant differences at *P* < 0.05 (one-way ANOVA and Tukey’s multiple comparison tests). Three biologically independent experiments with different batches of oocytes were carried out with similar results. (F) Structural analysis of Cys65 residue of ETR1. Left and Right, Cys65 and Tyr32 residues are shown in sticks in the ETR1 N-terminal AlphaFold structure with the side view and top view, respectively. (G) AlphaFold3 structural simulation of ETR1^C65Y^ mutation that induces steric clash.

The genetically identified ethylene receptor mutant *etr1-1* carries a single amino acid mutation C65Y, resulting in gain-of-function of ETR1 in *Arabidopsis* (Chang *et al*., 1993). Interestingly, according to the AlphaFold3 structural prediction, the C65 residue is located at the ion selective filter region of ETR1 N-terminal transmembrane domain (Figure 4F). Sequence alignments show that C65 is highly conserved in all ethylene receptors (Figure 4H). Thus, we simulated ETR^C65Y^ structure using AlphaFold3 and found that C65Y substitution introduces steric clash in the filter region (Figure 4G), which may block ion permeability. To verify the structural finding, we expressed ETR1^C65Y^ in *Xenopus oocytes* (Supplemental Figure 5B). Indeed, the TEVC recordings detected no Ca^2+^ inward currents, suggesting that the channel activity was completely blocked in ETR1^C65Y^ (Figure 4C-E). These findings support that Ca^2+^ permeable channel activity is highly relevant for ETR1 function.

### Ethylene evokes cytosolic Ca^2+^ influx that exclusively depends on subfamily I receptors

Based on the findings above, we next investigated whether ethylene regulates cytosolic Ca^2+^ levels. We first treated transgenic plants expressing the luminescent Ca^2+^ reporter aequorin (*AEQ*) (Ma et al., 2019) with ethylene and monitored chemiluminescence. Following ethylene treatment, we indeed observed a significant increase in cytosolic Ca^2+^ level (Figure 5A). To examine whether the ethylene-elicited Ca^2+^ influx is dependent on the subfamily I ethylene receptors, we used transient expression of Ca^2+^ reporter aequorin in the protoplasts derived from wild-type plant and various receptor mutants. Ethylene induced a rapid and transient luminescence spike in wild-type protoplasts, confirming an ethylene-triggered cytosolic Ca^2+^ influx (Figure 5B). Notably, protoplasts lacking subfamily I receptors (*iEIN2/11ein2*) exhibited no detectable Ca^2+^ increase upon ethylene treatment, in contrast to protoplasts from *iEIN2/ein2* and *iEIN2/224ein2* plants (Figure 5C). These results indicate that ethylene-induced cytosolic Ca^2+^ elevation requires functional subfamily I receptors, but not subfamily II receptors, consistent with the electrophysiological findings on distinct Ca^2+^-permeability of two subfamily receptors. Altogether, our data provide *in vivo* evidence for functional divergence of two ethylene receptor subfamilies in Ca^2+^-permeable channel activities.

**Figure 5.**
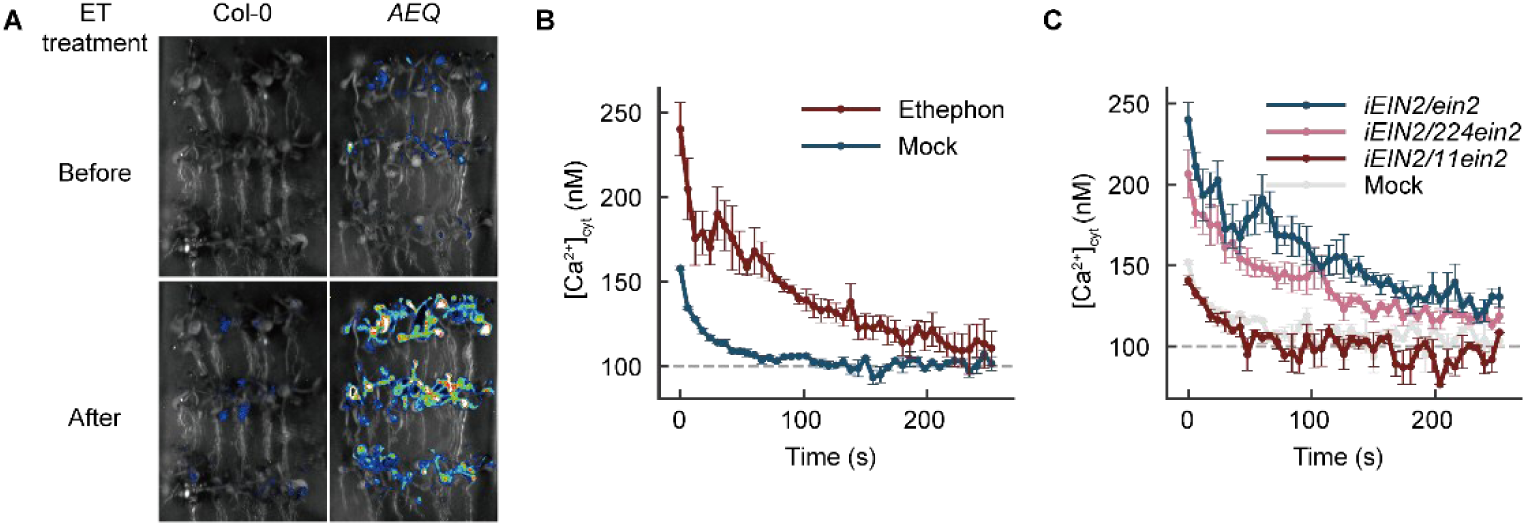
Ethylene elevates cytosolic Ca^2+^ level that exclusively depends on subfamily I receptors. (A) Images showing ethylene-induced changes in cytosolic Ca^2+^ concentration ([Ca^2+^]_cyt_) in 12-day-old wild-type and AEQ-expressing transgenic seedlings. (B) Time course analysis of 1.25 mM ethephon stimulated [Ca^2+^]_cyt_ elevation. For the mock control, the solvent of ethephon, i.e. the WI-Ca protoplast culture buffer, was used. Values represent means ± SE (n≥3). (C) Time course analysis of 1.25 mM ethephon stimulated [Ca^2+^]_cyt_ elevation in different genotypes. During the substrate incubation step, 1 μM β-estradiol was included to induce *EIN2* expression. For the mock control, the solvent of ethephon, i.e. the WI-Ca protoplast culture buffer, was used. Values represent means ± SE (n≥3).

## Discussion

Ethylene is a crucial phytohormone involved in the regulation of plant growth, development and responses to biotic and abiotic stresses (Schaller and Kieber, 2002). Over nearly four decades, substantial progress has been made in the identification of key signaling components and the elucidation of the ethylene signaling cascade (Hao *et al*., 2025). Although ethylene receptor ETR1 was the first plant hormone receptor identified genetically, the biochemical function of ethylene receptors remains elusive. Previous reports have shown that loss-of-function mutants of subfamily I and subfamily II ethylene receptors exhibit distinct phenotypes, implying functional divergence between the two receptor subfamilies (Hua and Meyerowitz, 1998; Qu *et al*., 2007). However, due to the extremely constitutive triple-response and postembryonic lethality, it was difficult for detailed phenotypic and molecular analyses of subfamily I double loss-of-function mutants (Qu *et al*., 2007). By using a tailored genetic strategy, we unequivocally demonstrated that ethylene perception was completely abolished in plants carrying double loss-of-function mutations in subfamily I receptors. Our results further indicate two subfamilies of receptors are not genetically in parallel, in which subfamily I receptors are epistatic to subfamily II receptors. Interestingly, subfamily II receptors are required for ethylene response mainly in relatively high dosage range of ethylene, as triple loss-of-function mutations in subfamily II receptors enable plants to only respond to low doses of ethylene. Although the underlying mechanism is currently unknown, it nicely aligns with evolutionary patterns as ACC OXIDASE (ACO), the ethylene-forming enzyme, and subfamily II receptors concurrently emerged and expanded in seed plants (Gallie, 2015; Li et al., 2018), hinting that the emergence of subfamily II receptors may represent an adaptive strategy enabling plants to cope with increased range of ethylene concentrations in seed plants.

Ethylene receptors contain a conserved N-terminal transmembrane domain. Through AlphaFold3-based structural simulations and electrophysiology assays, we discovered that subfamily I ethylene receptors function as Ca^2+^-permeable channels, while subfamily II receptors do not possess Ca^2+^-permeable activity. Ethylene receptors are known to be located in the ER (Chen *et al*., 2002), which is the main organelle for intracellular Ca^2+^ storage (Luan and Wang, 2021). Identification of Ca^2+^-permeable channel activity provides a possible explanation for the ER location of ethylene receptors. The ethylene receptor homodimers have been shown to be essential for ethylene perception and signaling transduction, the AlphaFold3-based structural predictions further reveal that homodimer of ethylene receptors is crucial for forming symmetrical ion channel-like structures. The findings that subfamily I receptors possess Ca^2+^-permeable activity and play predominant genetical roles in ethylene signaling suggest that Ca^2+^-permeable activity is highly relevant to the biological function of ethylene receptors. In support of this, genetically identified point mutation in *etr1-1* disrupt Ca^2+^-permeable activity of ETR1.

Both calcium and ethylene are known as important second messenger and phytohormone for plant response to biotic and abiotic stresses (Waadt et al., 2022; Zhai et al., 2022). Our finding that ethylene receptors function as calcium-permeable channels establishes a link between ethylene and calcium signaling, which is consistent with previous findings that ethylene could induce calcium influx in plant cells (Gravino et al., 2015; Hiraki et al., 2020). Interestingly, recent studies have uncovered previously unrecognized biochemical functions of some well-studied signaling components in triggering calcium signaling. For instance, plant disease resistance proteins were reported to form resistosomes functioning as calcium-permeable channels to activate plant immunity (Bi et al., 2021; Wang et al., 2019). Although the biological role of ethylene receptor-mediated calcium influx remains limited so far, the discovery of calcium-permeable channel activity of ethylene receptors provides a new perspective on ethylene signaling. Whether such calcium influx has an effect on the functionality of the C-terminal kinase domains of ethylene receptors, as well as downstream components such as CONSTITUTIVE TRIPLE RESPONSE 1 (CTR1) (Kieber et al., 1993) and EIN2, requires further investigation.

## Methods

### Plant materials and growth conditions

*Arabidopsis thaliana* ecotype Columbia-0 (Col-0) was used as the wild type in this study. The *etr1-7 ers1-3^+/−^* (Qu *et al*., 2007) and *etr2-3 ers2-3 ein4-4* (Hua and Meyerowitz, 1998) mutants are in a heterozygous Col-0/Ws (Wassilewskija) background and have been described previously. The *ers1-1* transgenic line is in the No-0 (Nossen) background (Hua *et al*., 1995), the *UBQ10::AEQ* (Ma *et al*., 2019) and *iEIN2-CFP-HA/ein2-5* (Li et al., 2015) transgenic lines are in the Col-0 background; all of these lines were reported previously. The *etr2-1* (Sakai *et al*., 1998) and *ein4-1* (Hua *et al*., 1998) mutants are also in the Col-0 background and have been described previously. Inducible *EIN2* lines in higher-order receptor mutant backgrounds, including *iEIN2-CFP-HA/etr1-7 ers1-3 ein2-5* (*iEIN2/11ein2*), *iEIN2-CFP-HA/etr2-3 ers2-3 ein4-4 ein2-5* (*iEIN2/224ein2*), and *iEIN2-CFP-HA/etr1-7 ers1-3 etr2-3 ers2-3 ein4-4 ein2-5* (*iEIN2/11224ein2* were generated in this study by genetic crossing. *ers1-1 etr1-7 ers1-3*, *ein4-1 etr1-7^+/-^ ers1-3* and *etr2-1 etr1-7^+/-^ ers1-3* were also generated in this study by genetic crossing.

Seeds were surface-sterilized in 75% (v/v) ethanol containing 0.02% (v/v) Triton X-100, and then plated on MS medium (supplemented with 1% sucrose and 0.8% agar) or on half-strength MS (½ MS) medium (supplemented with 0.8% agar). The plates were stratified at 4 °C for 3–4 days.

For triple response assays, plates were exposed to light for 4 h after stratification at 4 °C and subsequently incubated in the dark for 3.5 days. For all other experiments, plates were transferred after stratification to a growth room at 22 °C under a 16 h light/8 h dark photoperiod, with a photosynthetic photon flux density of 60 μmol m⁻² s⁻¹ and a color temperature of 4500 K.

For ethylene treatments, seedlings were exposed to 10 p.p.m. ethylene. For 1-MCP treatment, a supersaturated concentration (≥100 p.p.m.) was used.

For the dynamic platform assay, surface-sterilized seeds were stratified at 4 °C for 3 days and then spotted onto MS medium in square Petri dishes. Using a stereomicroscope and a sterile syringe needle, the seeds were gently oriented with the micropyle facing downward. The plates were exposed to white light on a dynamic imaging platform (Dynaplant®, Beijing MICROLENS) for 4 h, after which the white light was switched off and images were acquired every 2 h for a total of 96 h. For each seedling, the time of germination was recorded, and the image taken 48 h after germination was used to measure hypocotyl length using ImageJ software (https://imagej.nih.gov/ij/). The measurements were then compiled for statistical analysis.

### Genotyping

The *ers1-3, ers2-3, and the iEIN2-CFP-HA* T-DNA insertion lines were identified by PCR-based genotyping. For *ers1-3,* the primer pair **e**rs1-3LP/ers1-3RP was used to amplify the wild-type reaction (WR), and ers1-3LP/pSKI015-LB was used for the T-DNA reaction (TR). For *ers2-3*, J2/J7 was used as the WR primer pair and jLB3/J2 as the TR primer pair. For the *iEIN2-CFP-HA* insertion, iE2JL-F/iE2JL-R served as the WR primer pair, and pER8-LB3/iE2JL-R as the TR primer pair. Primer sequences are listed in the supplemental table. *ein2-5*, *etr1-7*, *etr2-1*, *etr2-3*, *ein4-1*, and *ein4-4* were genotyped using derived cleaved amplified polymorphic (dCAPS) markers (Neff et al., 1998). The corresponding PCR products were digested with Bsp119I, EcoRI, XmaJI, HinfI, TaaI, or MnlI (Thermo Scientific), respectively, and separated on 2.5–4% agarose gels by electrophoresis. For all dCAPS markers, wild-type PCR products were designed to contain a restriction site and thus could be cleaved by the enzyme, whereas mutant PCR products remained undigested. dCAPS marker primer sequences are provided in supplemental table.

### Hypocotyl and root measurements

Images of seedlings were captured with a digital camera (Canon, EOS 760D). Hypocotyl and root lengths were then measured for at least 20 and 10 seedlings, respectively, using ImageJ software.

### Total protein extraction and western blot analysis

*Arabidopsis* seedlings were ground to a fine powder in liquid nitrogen and homogenized in an equal volume of 4× SDS sample buffer (250 mM Tris-HCl, pH 6.8, 40% (v/v) glycerol, 8% (w/v) SDS, 0.05% (w/v) bromophenol blue, 50 mM DTT [freshly added], and 0.04% (v/v) β-mercaptoethanol [freshly added]). The extracts were mixed thoroughly for 5–10 min and then heated at 65 °C for 10 min. After centrifugation at 20,000 × g for 5 min, the supernatants were separated by 7.5% SDS–PAGE and transferred to PVDF membranes for immunoblotting. Immunoblot analyses were performed using specific primary antibodies, including anti-EIN3 (Guo and Ecker, 2003) and anti-HSP90 (Beijing Protein Innovation), together with HRP-conjugated secondary antibodies (anti-rabbit IgG–HRP and anti-mouse IgG–HRP; Promega). Signal intensities were quantified using ImageJ software.

### Total RNA extraction and real time qPCR

Total RNA was extracted from *Arabidopsis* seedlings using the Eastep^®^ Super Total RNA Extraction Kit (Promega). First-strand cDNA was synthesized from total RNA using the All-in-one^®^ 5× RT MasterMix (abm). Quantitative real-time PCR (qPCR) was performed to determine transcript levels with the Hieff UNICON^®^ Universal Blue qPCR SYBR Green Master Mix (Yeasen). *PP2AA3* was used as the reference gene. SE values were calculated from at least three replicates. Primer sequences are provided in the supplemental table.

### Transcription in vitro

To construct *Xenopus oocyte* expression vectors, an intermediate plasmid (pT7Ts-eGFP) was first generated by inserting the eGFP coding sequence, amplified from the pMBP-GFP vector, into the pT7Ts backbone using Gibson assembly. The coding sequences (CDS) of *ETR1*, *ETR1^C65Y^*, *ERS1*, *ETR2*, *ERS2*, and *EIN4* were then amplified from cDNA and cloned into pT7Ts-eGFP (linearized by PCR to two fragments: BB-pT7Ts and lk-eGFP-pT7Ts) by multiple fragment homologous recombination. A short flexible linker (NIGSGSNGSSGS) was inserted between each receptor CDS and eGFP to minimize any interference of the fluorescent tag with receptor function. The point mutation F76Y in *ETR1* (*ETR1^F76Y^*) was introduced by site-directed mutagenesis using the pT7Ts-ETR1-eGFP construct as a template. A list of primers used for vector construction is provided in Table.

To obtain the transcription template, plasmid DNA was first prepared using the Plasmid Maxprep Kit (Vigorous). The purified plasmid was linearized with KpnI, and the linearized fragment was recovered. All subsequent steps were performed under RNase-free conditions. The linearized DNA was treated with Proteinase K to digest RNases and then recovered. In vitro transcription was carried out using the mMESSAGE mMACHINE T7 Kit (Invitrogen), followed by DNase digestion to remove the DNA template and subsequent recovery of the cRNA. The quality of the linearized DNA fragment, Proteinase K-treated product, in vitro transcription product, DNase digestion product, and final cRNA was assessed by agarose gel electrophoresis and Nanodrop.

### Two-electrode voltage clamping (TEVC) recording from *Xenopus oocytes*

TEVC recording assays were performed as described (Ming et al., 2025). The coding sequences for target genes were inserted into the pT7Ts vector. cRNAs were in vitro transcribed from the linearized pT7Ts vectors using the mMESSAGE mMACHINE T7 kit (Invitrogen). cRNA quality was verified by Nanodrop, and the oocytes were injected with 25 or 50 ng cRNAs or distilled water. The injected oocytes were incubated at 18 °C for 48 h in ND96 [96 mM NaCl, 2 mM KCl, 1 mM MgCl_2_, 1.8 mM CaCl_2_, 5 mM HEPES-KOH (pH 7.5)] supplemented with 0.05 mg/mL gentamycin and 0.1 mg/mL streptomycin. The protein expression was detected by using eGFP fluorescence signals with a Leica STELLARIS5 confocal microscope. eGFP signals were detected with excitation at 488 nm. TEVC recordings were performed using a bath solution containing 30 mM CaCl₂, 2 mM NaCl, 1 mM KCl, 130 mM mannitol, and 5 mM MES-Tris, pH 5.6. Measurements were conducted using an Axoclamp 900 A Computer-Controlled Microelectrode Amplifier (Axon Instruments). Voltage steps ranged from +40 mV to -140 mV in -20 mV decrements (2 sec duration). Each step began with 0.03 sec and ends with 1.2 sec at the resting potential of the oocyte membrane in the recording solution. Currents were digitized via a Digidata 1440A converter and recorded with Clampex 10.7 software (Axon Instruments), using a pipette solution of 3 M KCl. TEVC experiments were conducted using a minimum of three distinct batches of oocytes, sourced from different frogs and prepared on separate days.

### Aequorin-based calcium signal imaging

*UBQ::AEQ* seedlings were grown on ½ MS medium for 12 days. Seedlings were then uniformly sprayed with 50 μM coelenterazine h (NanoLight Technologies) and incubated in the dark for 12 h. Thereafter, the plants were transferred onto fresh medium, sealed in resealable plastic bags, and aequorin bioluminescence images were recorded. Following this initial acquisition, ethylene was applied inside the sealed bags, and bioluminescence images were recorded again using the same exposure time as in the first recording.

### Aequorin-based cytosolic Ca^2+^ measurements

Plasmid DNA was first prepared using the Plasmid Maxprep Kit (Cwbio). Protoplasts from different genotypes were isolated and transformed as previously described (Yoo et al., 2007). The culture medium was supplemented with 2 mM CaCl₂ (WI-Ca) according to established protocols (Maintz et al., 2014). After overnight incubation for 10-12 h, protoplasts were collected and incubated with 10 μM coelenterazine h for 1-4 h. They were then centrifuged to remove the coelenterazine h containing medium and resuspended in fresh WI-Ca buffer, adjusting the cell density to the same level per milliliter. Protoplast suspensions were dispensed into white 96-well plates and subjected to treatments in the following order: WI-Ca (mock), 3 mM H₂O₂ (to determine total luminescence capacity), and 1.25 mM ethephon, so as to avoid interference from ethylene released by ethephon. For each condition, 3-6 wells were measured in parallel. Luminescence was recorded every 6 s with an integration time of 0.5 s per well. Stimulus (or solvent for mock) was manually added at t = 0, and signals were collected for 600 s (plots truncated to 252 s where indicated). Each condition had 3-4 technical replicates. Data are shown as mean ± SE. Peak amplitude and peak time were extracted from the time course. T1/2 was defined as the time from the peak to halfway toward the 100 nM baseline. AUC(0-252 s) was computed using the trapezoidal rule. Unless stated otherwise, all statistics were performed on the unscaled [Ca^2+^]_raw_ traces; time points were masked once L_remain_(t) fell below 5%. For each run, a shared L_total_ was estimated from slow-peak wells elicited by 3 mM H_2_O_2_ and applied across all wells of the same experiment. Traces were then transformed with the Allen/Hill procedure and K-anchored by setting the mock tail (last 20 points) to 100 nM, providing a stable and comparable nM scale without absolute photon calibration (Allen et al., 1977).

### Statistical analysis

Statistical analyses were performed using one-way, two-way or three-way ANOVA followed by post hoc multiple-comparison tests. Significant differences between samples were indicated by different lowercase letters for *P* ≤ 0.05 and by different uppercase letters for *P* ≤ 0.01, or by directly reporting the corresponding *P* values.

## Author contributions

H.W.G., W.S., and L.L. conceived the project; H.W.G., C.L.P., W.S., and L.L. designed the experiments; C.L.P. generated the genetic materials, collected the genetic phenotype and performed western blot assays, qRT-PCR assays, and AEQ-based [Ca^2+^]_cyt_ assays; Z.N.X. performed large-scale plasmid preparations for AEQ-based [Ca^2+^]_cyt_ assays in protoplasts; J.Y.C. and Z.N.L performed the electrophysiology assays in *Xenopus Oocyte* and structural analysis with assistant of Y.H.M.; C.L.P., W.S., D.D.H., L.L. and H.W.G. wrote the manuscript.

## Acknowledgments

We thank Dr. Yan Guo (China Agricultural University) for kindly providing *UBQ10::AEQ* seeds; We thank the microscopy and electrophysiological facilities of State Key Laboratory of Plant Environmental Resilience in China Agricultural University. We also thank all members in the H.G. laboratory for helpful discussions and suggestions. This work was supported by the funds from National Natural Science Foundation of China (32230008 and W2521010 to H.G.; 32300275 to D.H.; 32470301 to L.L.), the New Cornerstone Science Foundation (Grant No. NCI202235 to H.G.), China Agricultural University Fund (2025RC042 to W.S.), State Key Laboratory of Plant Environmental Resilience Fund (SKLPERKF2504 to W.S.), the Pinduoduo-China Agricultural University Research Fund (PC2024B01003 to W.S.), and Shenzhen Science and Technology Program (Grant No. KQTD20190929173906742, ZDSYS20230626091659010 to H.G.).

---

**Supplemental Figure 1.**
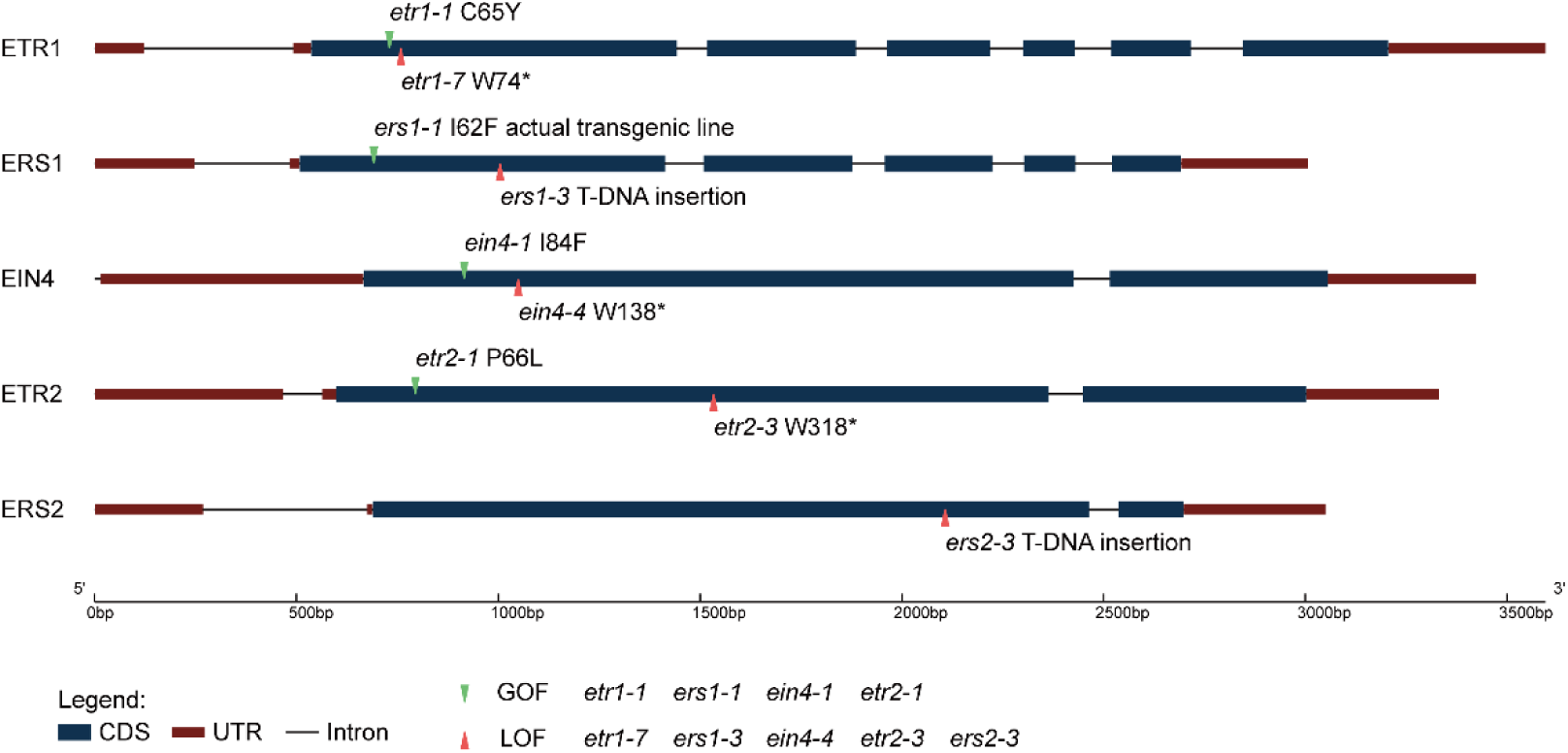
Schematic overview of five ethylene receptors mutant alleles in *Arabidopsis*. Gain-of-function mutants are indicated in green and labeled above the corresponding genes, whereas loss-of-function mutants are indicated in red and labeled below the genes. For *ers1-1*, the mutant *ERS1* gene was introduced into different genetic backgrounds by transgenic approach.

**Supplemental Figure 2.**
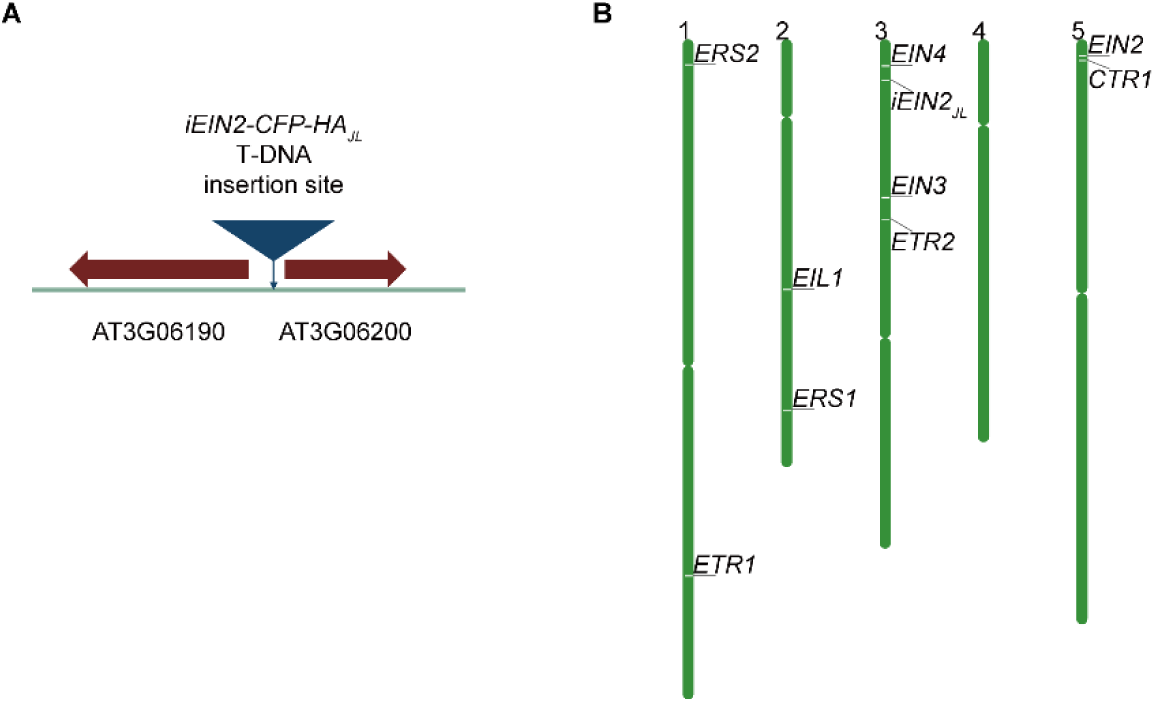
Schematic genomic locations of the ethylene receptor genes. (A) Schematic representation of the insertion site of the *iEIN2-CFP-HA* transgene expression cassette. The transgene is inserted in the proximal region of chromosome 3, between AT3G06190 and AT3G06200. (B) Schematic representation of the genomic locations of the genes analyzed in this study. The insertion site of the *iEIN2-CFP-HA* transgene is tightly linked to *EIN4*.

**Supplemental Figure 3.**
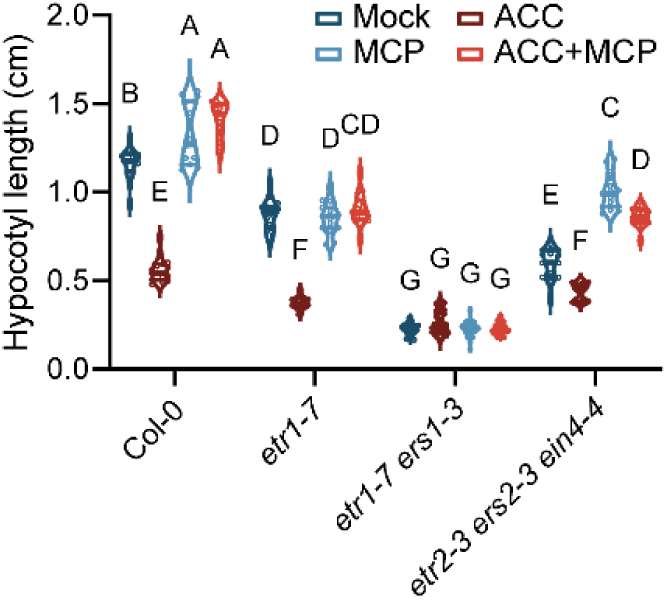
1-MCP treatment suppresses the loss-of-function mutants of subfamily II but not subfamily. **I.** Quantification of hypocotyl length in 3.5-day-old etiolated seedlings on MS medium under four treatments: Mock, ACC (20 μM), MCP, and ACC (20 μM)+MCP. 1-MCP (gaseous) as supplied by slow release to a saturated concentration (>100 p.p.m.) in sealed containers. Statistical significances between samples were analyzed by three-way ANOVA with multiple comparisons and indicated by different uppercase letters for *P*≤0.01 (n≥14).

**Supplemental Figure 4.**
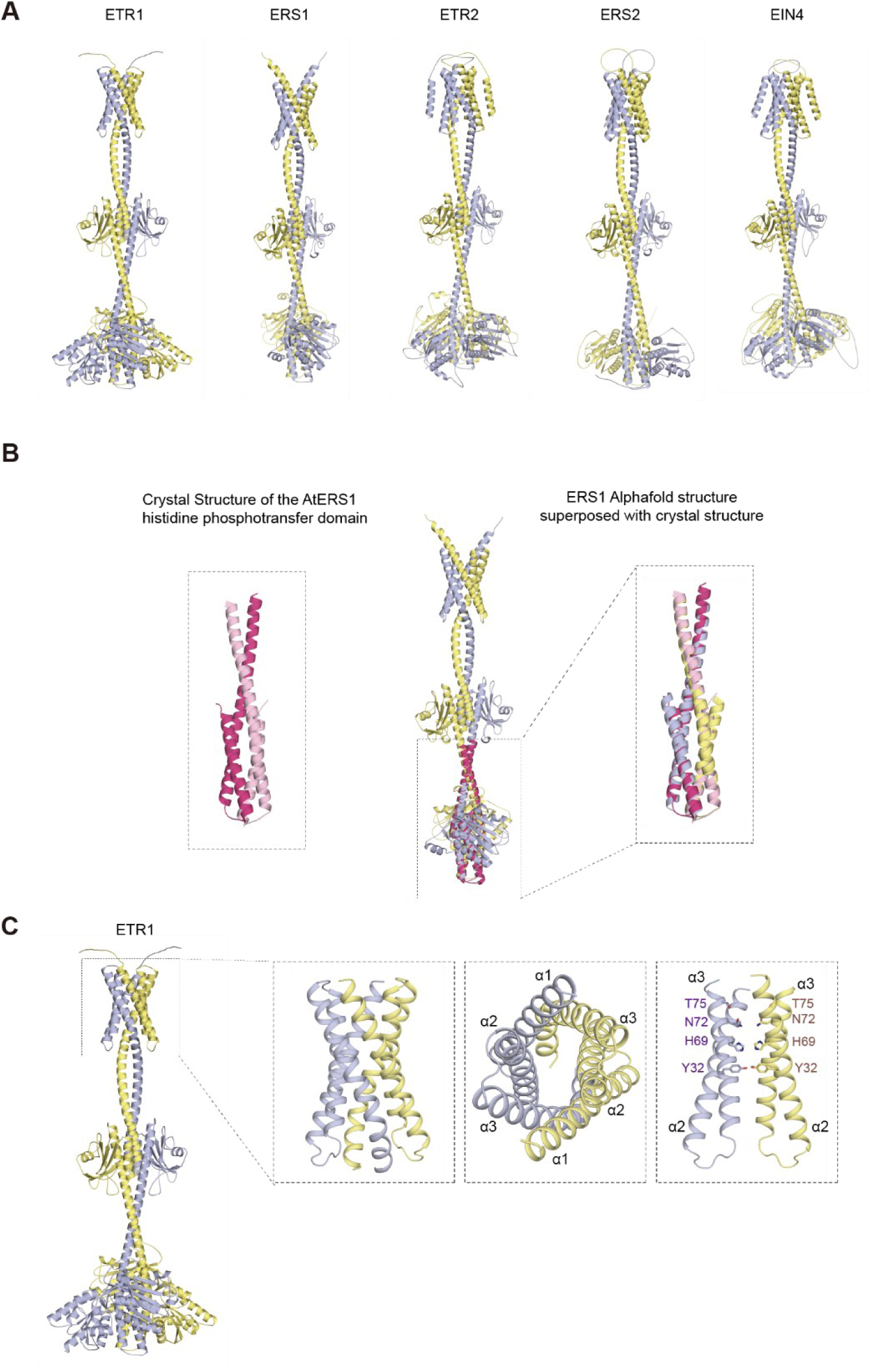
AlphaFold3 structural simulations of ethylene receptors. (A) The AlphaFold3 structural simulations of ethylene receptor homodimers ETR1, ERS1, ETR2, ERS2, EIN4. Ethylene receptor protomers are shown in colors of light blue and yellow. (B) The structural superposition of ERS1. The superposition of AlphaFold3 structure and crystal structure of ERS1 histidine phosphotransfer domain (PDB:4MT8) are highlighted. (C) The AlphaFold3 structural simulations of ethylene receptor ETR1 homodimers. The N-terminal transmembrane domains are highlighted. The α-helices are labeled. The charged residues in the α-helix 2 and α-helix 3 are indicated.

**Supplemental Figure 5.**
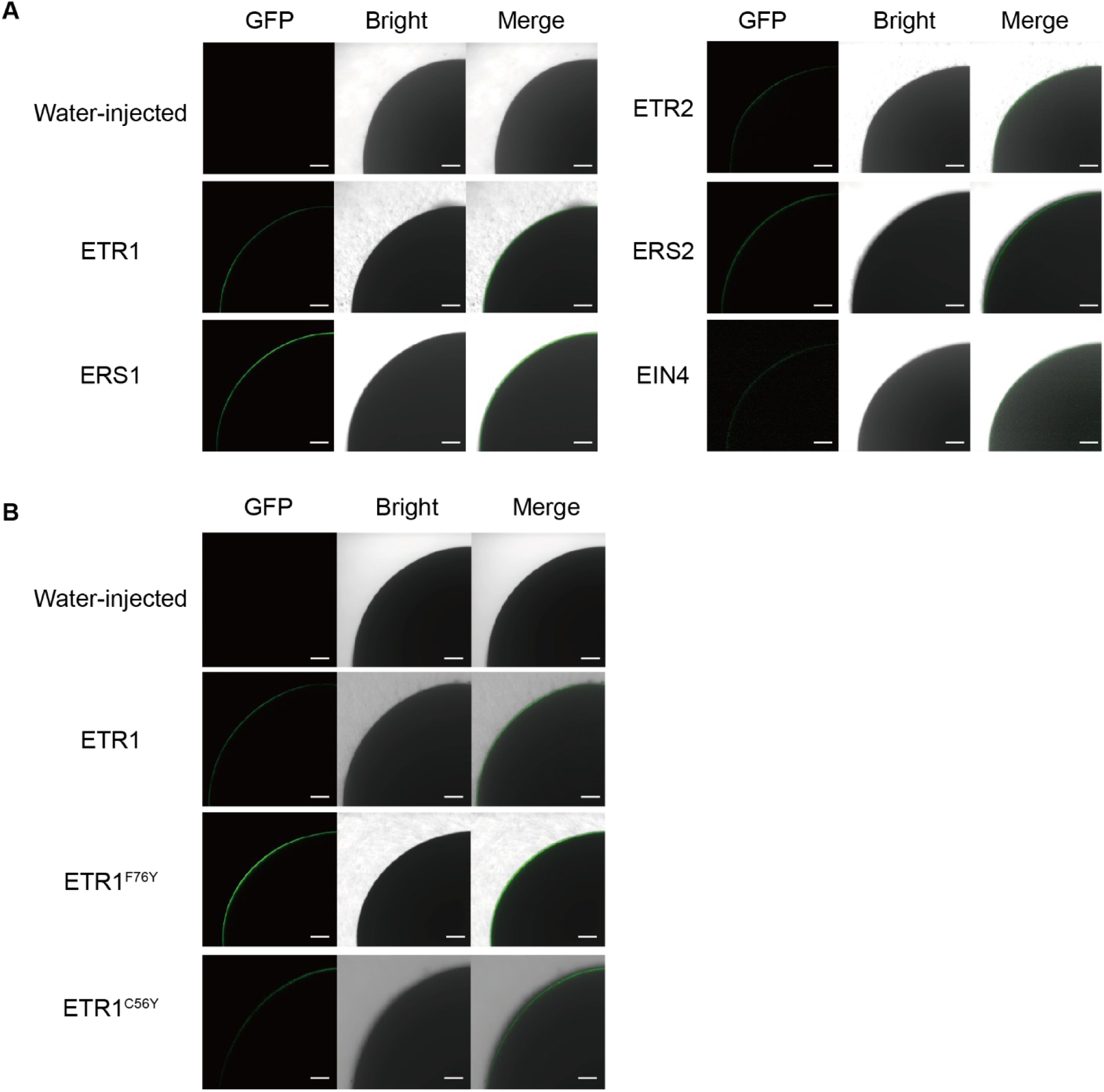
Confocal fluorescence images of *Xenopus oocytes* expressing wild-type or mutant ethylene receptors. (A) Confocal fluorescence images of *Xenopus oocytes* expressing ETR1-eGFP, ERS1-eGFP, ETR2-eGFP, ERS2-eGFP or EIN4-eGFP. eGFP signals were observed by using confocal microscopy. *Xenopus oocytes* injected with water serve as a negative control. Scale bars, 100 μm. (B) Confocal fluorescence images of *Xenopus oocytes* expressing ETR1-eGFP or its variants. eGFP signals were observed by using confocal microscopy. *Xenopus oocytes* injected with water serve as a negative control. Scale bars, 100 μm

**Supplemental Table 1.**
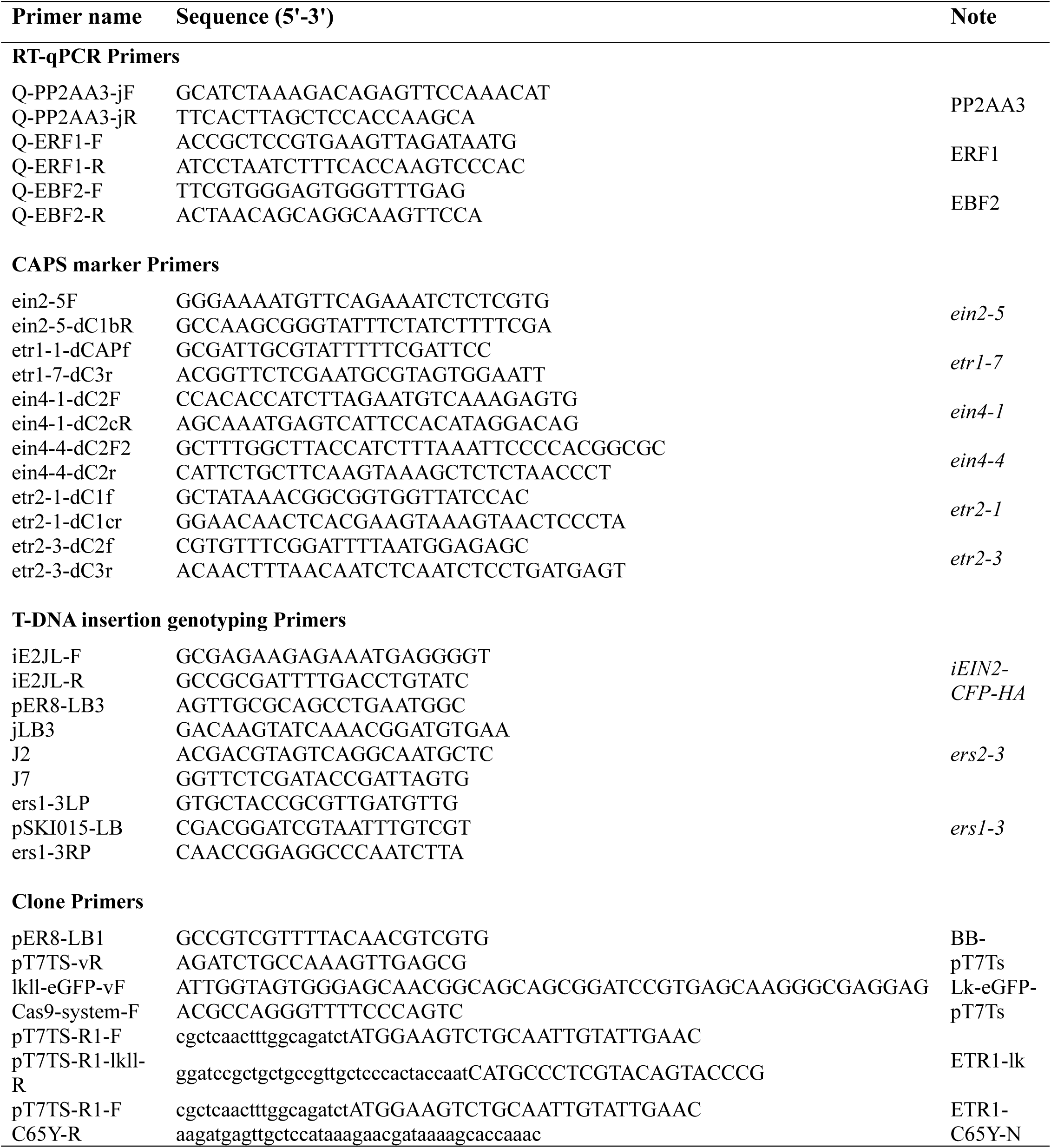
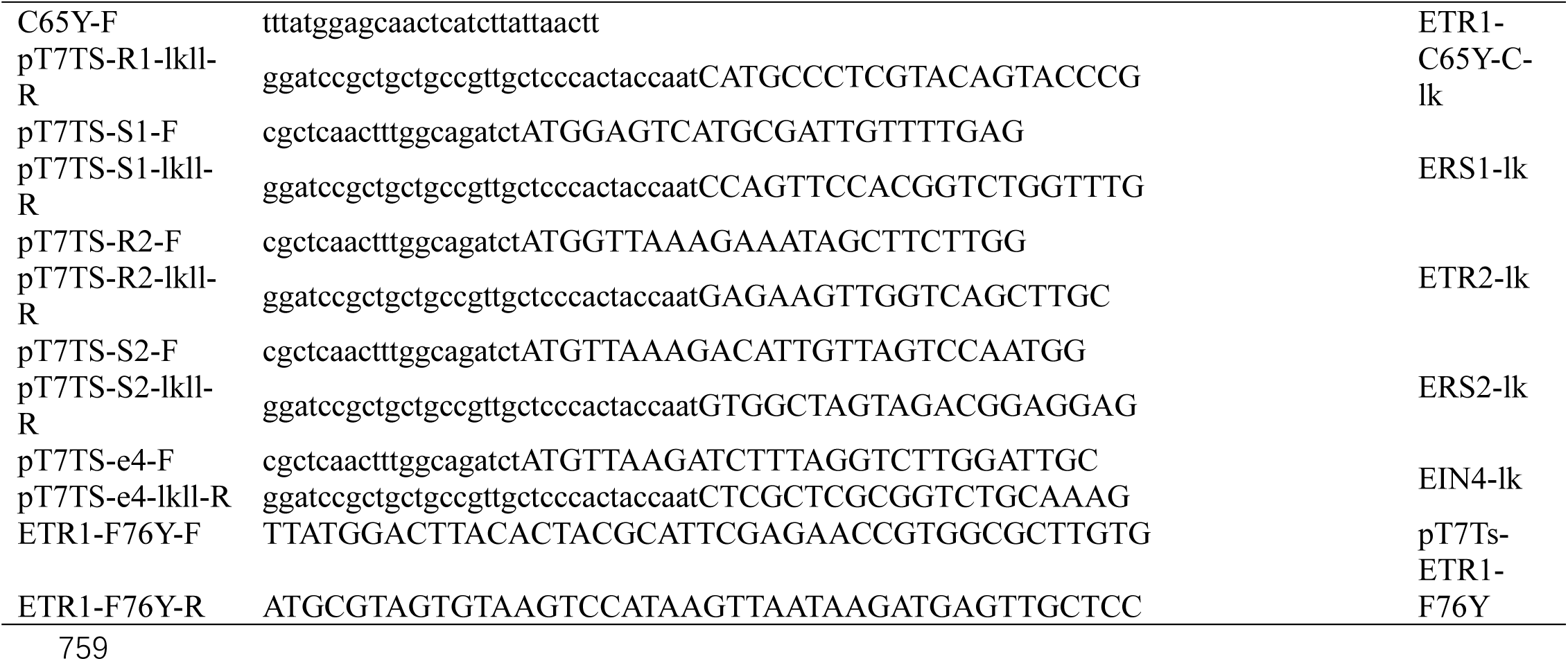
Primers used in this study.

